# Transcriptome Profiling of Mouse Brain and Lung under Dip2A Regulation Using RNA-Sequencing

**DOI:** 10.1101/563916

**Authors:** Rajiv kumar sah, Anlan Yang, Fatoumata Binta Bah, Salah Adlat, Ameer Ali Bohio, Zin Mar Oo, Chenhao Wang, May zun Zaw Myint, Noor Bahadar, Luqing Zhang, Xuechao Feng, Yaowu Zheng

**Author notes:** (LQZ), (XCF); (YWZ).

## Abstract

Disconnected interacting 2 homolog A (DIP2A) gene is highly expressed in nervous system and respiratory system of developing embryos. However, genes regulated by Dip2a in developing brain and lung have not been systemically studied. Transcriptome of brain and lung in embryonic 19.5 day (E19.5) were compared between wild type and Dip2a^-/-^ mice. Total RNAs were extracted from brain and lung of E19.5 embryos for RNA-Seq. Clean reads were mapped to mouse reference sequence (mm9) using Tophat and assembled into transcripts by Cufflinks. Edge R and DESeq were applied to identify differentially expressed genes (DEGs) and annotated under GO, COG, KEGG and TF. An average of 50 million reads per sample was mapped to the reference sequence. A total of 214 DEGs were detected in brain (82 up and 132 down) and 1900 DEGs in lung (1259 up and 641 down). GO enrichment analysis indicated that DEGs in both Brain and Lung were mainly enriched in biological processes ‘DNA-templated transcription and Transcription from RNA polymerase II promoter’, ‘multicellular organism development’, ‘cell differentiation’ and ‘apoptotic process’. In addition, COG classification showed that both were mostly involved in ‘Replication, Recombination and Repair’, ‘Signal transduction and mechanism’, ‘Translation, Ribosomal structure and Biogenesis’ and ‘Transcription’. KEGG enrichment analysis showed that brain was mainly enriched in ‘Thryoid cancer’ pathway whereas lung in ‘Complement and Coagulation Cascades’ pathway. Transcription factor (TF) annotation analysis identified Zinc finger domain containing (ZF) proteins were mostly regulated in lung and brain. Interestingly, study identified genes Skor2, Gpr3711, Runx1, Erbb3, Frmd7, Fut10, Sox11, Hapln1, Tfap2c and Plxnb3 from brain that play important roles in neuronal cell maturation, differentiation and survival; genes Hoxa5, Eya1, Ctsh, Erff1, Lama1, Lama2, Rspo2, Sox11, Spry4, Shh, Igf1 and Wnt7a from lung are important in lung development and morphogenesis. Expression levels of the candidate genes were validated by qRT-PCR. Genome wide transcriptional analysis using wild type and Dip2a knockout mice in brain and lung at embryonic day 19.5 (E19.5) provided a genetic basis of molecular function of these genes.

## Introduction

DIP2A is a member of Disconnected (*disco*)-interacting 2 (DIP2) protein family whose molecular anatomical function remains to be clarified. DIP2A was firstly identified in Drosophila as a novel transcription factor that interacts with Disconnected (*disco*) gene needed for proper neural connection during visual system development in Drosophila [1-3]. Previous studies have shown that Dip2A is highly expressed in human brain and may play a role in axon patterning in Central Nervous System (CNS) [4]. Bioinformatics analysis using Homolog Gene suggests that DIP2A is a receptor molecule with DMAP, AMP and CaiC binding domains [5]. At DNA replication site, DIP2A, in a complex with DMAP1-DNMT1-HDAC, regulates neurite outgrowth and synaptic plasticity [6]. Moreover, Dip2a has been previously identified as a risk gene associated with neurodevelopment diseases like autism spectrum disorder, Development dyslexia and Alzheimer diseases [7-9]. These all evidence strongly supports the role of Dip2a gene in both vertebrate and invertebrate nervous system development. However, which biological process or molecular function is regulated by Dip2a gene during embryonic brain development is not known.

In order to systematically understand expression pattern and physiological role of Dip2a gene, Dip2a-LacZ knockin and Dip2a 65kb knockout (Dip2a^-/-^) mouse models were generated using CRISPR/Cas9 system [10]. Dip2a expression via endogenous expression of β-Galactosidase gene (LacZ) have shown that Dip2a is highly expressed in brain neurons, retinal ganglia cell, reproductive, vascular and Lung tissue in adult and ectodermal tissue in developing embryos [11]. RNA sequencing (RNA-Seq) has rapidly emerged as a favorite approach for high throughput gene expression and function studies. Through RNA-Seq, gene expression and gene interactions at any time point or in a particular tissue can be investigated [12]. In present study, Transcriptome (RNA-seq) analysis of E19.5 brain and lung of WT and Dip2a^-/-^ embryo was performed.

Dip2a role in brain and lung development has not been studied before. A global Transcriptome analysis of brain and lung will help us in understanding of Dip2a function in regulating brain and lung development. A total of 214 genes in brain and 1900 genes in lung were identified differentially expressed under Dip2A, suggesting that these genes are potentially relevant to brain and lung development and function. Those genes are further explicated and discussed in this study.

## Materials and methods

### Animals

All procedures were conducted following guidelines recommended in the guide for Care and Use of Laboratory Animals of National Institutes of Health with approval of Institutional Animal Care and Use Committee of Northeast Normal University (NENU/IACUC, AP2013011). Mice were housed in clean facility in individual IVC cages under a normal 12:12h light:dark cycles in a temperature of 20°C and humidity 50 ± 20% in Northeast Normal University. All mice were anesthetized before euthanasia with 1% pentobarbital at a dose of 10mg/kg and all effort was made to minimize suffering.

### RNA isolation and library preparation for RNA-Seq

Dip2a heterozygous mice (Dip2a^+/-^) were intercrossed, female mice checked for presence of copulation plug (Vaginal plug) and designated as E0.5 day. At age of E19.5, pregnant dams were euthanized and embryos were collected on ice-cold 1X PBS. Brains and lungs were dissected out from each embryo and frozen immediately in liquid nitrogen. Total RNA from brain and lung was isolated using RNAiso plus reagent (Takara, Dalian) in accordance with the manufacturer’s instruction and followed by additional step of DNase I digestion to eliminate genomic DNA contamination. The quality and purity of RNA was checked by Nano drop ND-1000 spectrophotometer (Thermo Fisher Scientific, USA) and Agilent 2100 Bio analyzer (Santa Clara, CA, USA). Library was constructed and sequenced using Illumina Hiseq™ 2500 (Biomarker, Beijing, China).

### Sequence Mapping, assembly and gene annotation

Clean data (clean reads) were obtained by filtering out adapter only reads, ambiguous “N” base containing reads (N>5%) and reads with low Q-value (< 10) from raw data. The clean reads were then mapped to mouse reference genome (mm9) using Tophat v2.2.0 that allows up to two mismatches [13]. The mapped reads were then assembled into transcripts using Cufflinks v2.2.1 [13]. The transcripts were defined as unigenes and analyzed via BLASTX alignment (E-value < 1.0E^-5^) with NR (NCBI non-redundant protein) and Swiss-Prot databases under BLAST2GO platform [14-15].

### Gene expression quantification and analysis of differentially expressed genes (DEG)

Quantification of transcript expression levels was presented by FPKM (fragments per kilo base of exon per million fragments mapped) that minimize the reads output variations between samples. In order to identify differentially expressed genes (DEGs) between WT and Dip2a^-/-^ embryos in brain and lungs, we used DESeq software from R package [16]. Resulting P values were adjusted using the Benjamini and Hochberg’s approach for controlling the false discovery rate (FDR). Differentially expressed genes (DEGs) with a threshold FDR adjusted, p value<0.001 and fold change ≥ 2 (log2> ±1) were selected for further analysis. Gene Ontology (GO) enrichment analysis of DEGs was implemented by GOseq R packages [17]. KOBAS was used to test the statistical enrichment of DEGs in KEGG pathways. For transcription factor analysis, Genes were subjected under Animal TFDB database (Zhang et al., 2012).

### Quantitative real time PCR (qPCR) validation of RNAseq

One microgram of total RNA from brain and lung tissue of E19.5 WT and Dip2a^-/-^ embryos was reverse transcribed using primescriptTMII cDNA synthesis kit (Takara, Dalian, China). QPCR was performed using Thermo cycler (AnalytikJena AG, Jena, Germany) and SYBR II premix (Takara, Dalian, China). All results were normalized to housekeeping gene 18S ribosomal RNA and relative quantification was calculated using comparative threshold cycle (2^-ΔΔCt^) values for 3 biological replicates.

## Results and Discussion

### Gene expression profiling of brain and lung from WT and Dip2a^-/-^ mice

Four cDNA libraries were prepared from brain and lung of WT and Dip2a^-/-^ E19.5 embryos (n=3 per sample) and sequenced using Illumina HiSeq2000. After filtering out adaptors sequence and low quality reads, 24.08 GB of Clean Data were obtained, or 6.02 GB per sample, with a Q30 base percentage above 92.49%. The clean reads from each sample were then mapped to mouse reference genome (mm9) and quantification of transcripts expression levels were calculated and presented by FPKM. As shown in Table 1, the matching efficiency between the clean read and the reference genome of each sample ranged from 89.25% to 91.95%. On an average, about 6000 unigenes were expressed in each sample. Unigenes comparison between WT and Dip2a^-/-^ identified 5787 unigenes overlap in all sample and only 2 and 4 unigenes unique in WT brain and WT lung respectively (Fig 1).

**Table 1.** Statistics of sequence output

| Sample | Total clean reads | Total mapped reads | Q30 % | GC % |
| --- | --- | --- | --- | --- |
| WT brain | 40,288,890 | 37,047,078 (91.95%) | 96.99 | 46.12 |
| WT lung | 54,035,446 | 48,953,863 (90.60%) | 96.87 | 47.35 |
| Dip2a<sup>-/-</sup> brain | 52,633,352 | 47,472,439 (90.19%) | 96.78 | 46.89 |
| Dip2a<sup>-/-</sup> lung | 54,634,002 | 48,763,438 (89.25%) | 96.8 | 47.97 |

**Fig 1.**
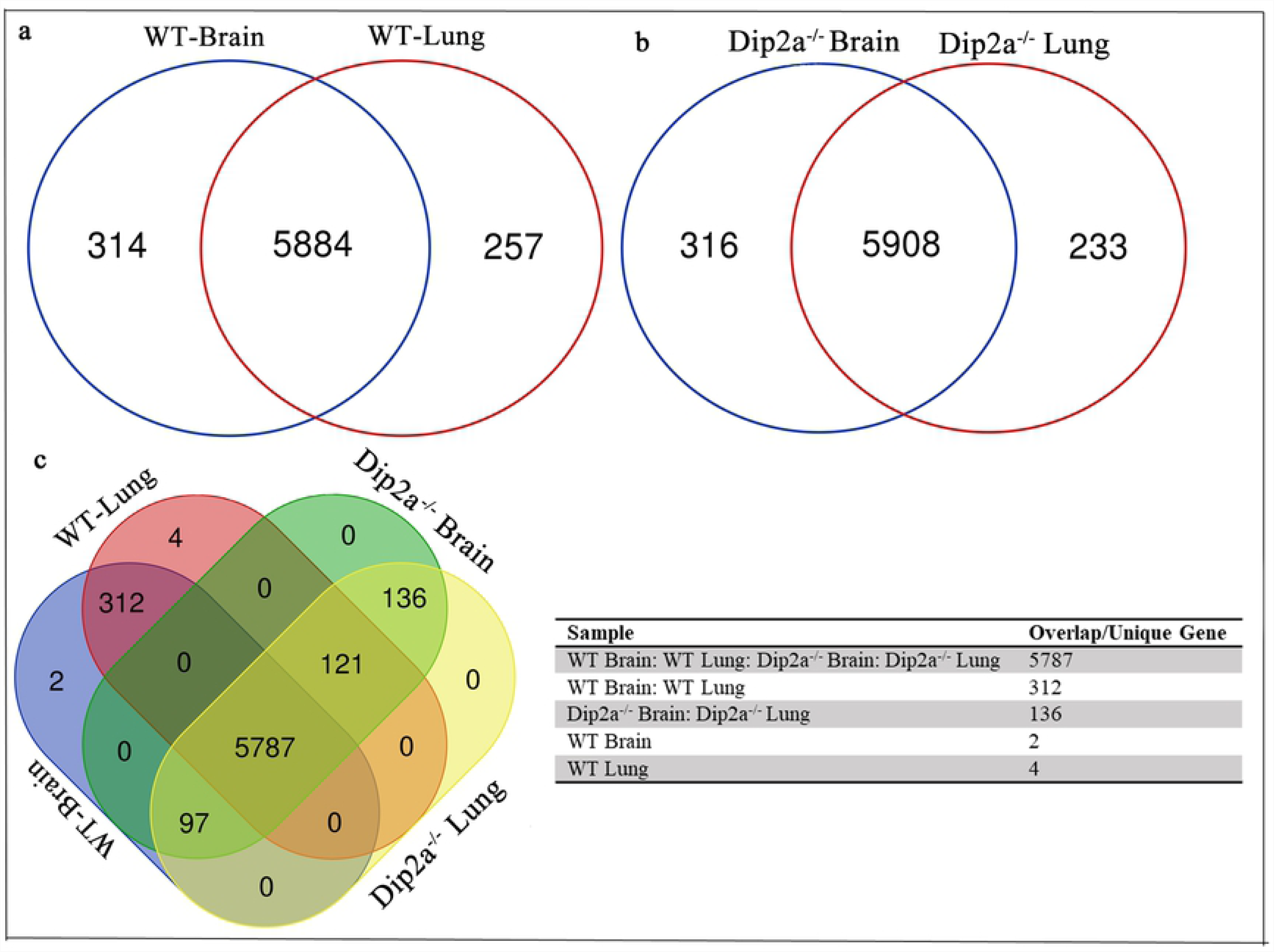
Venn diagrams showing overlap and unique unigenes identified within Wild type (WT) brain, Wild type (WT) lung, Dip2a^-/-^ brain and Dip2a^-/-^ lung. (a) In WT group, total 5884 unigenes were expressed in both samples, 314 unigenes unique to brain and 257 unigenes unique to lung (b) In Dip2a^-/-^ group, total 5908 unigenes were expressed in both samples, 319 unigenes were unique to brain and 233 unigenes were unique to lung. (C) 5787 unigenes were expression in all samples, 2 unigenes were unique to WT brain and 4 unigenes were unique to WT lung.

### Identification of differentially expressed genes and functional annotation

To identify differentially expressed genes, unigenes from WT brain vs. Dip2a^-/-^ brain and WT lung vs. Dip2a^-/-^ lung were compared. Cuffdiff identified 214 genes in brain and 1900 genes in lung to be differentially expressed, with Fold Change ≥2 and FDR < 0.01. In Dip2a^-/-^ brain, 82 genes were up-regulated and 132 genes were down-regulated whereas in Dip2a^-/-^ lung, 1259 genes were up-regulated and 641 genes were down-regulated when compared to WT (Fig 2). In Dip2a^-/-^ brain, Rpsa-ps10, Tpm3-rs7, Amd2 and Gm8730, Gm10709, Gm6768 and Gm9825 genes were highly over expressed whereas Acp5, Ifi204, Col10a1, Ibsp and Mmp13 genes were highly under expressed. Similarly, in Dip2a^-/-^ lung, genes like Rps2-ps6, Gm10709, Bhmt and Gm8730 were highly increased whereas genes like Il1r2, Nr4a3, Cela1 and Dlk2 were significantly decreased (Table 2). Functional annotation of brain and lung DEGs shows that more than 90 % of DEGs from brain and lung had significant matches in Nr, EggNOG, GO, COG, KEGG and Swiss-Prot database respectively (S1 Table).

**Table 2.** Highly significant differentially expressed genes in Dip2a^-/-^ group compared to Wild type group (FDR< 0.01, FC> 20)

| Gene ID | Gene symbol | WT Brain | Dip2a<sup>-/-</sup> Brain | FDR | log2FC |
| --- | --- | --- | --- | --- | --- |
|  |  | FKPM | FKPM |  |  |
| ENSMUSG00000047676 | Rpsa-ps10 | 0.91 | 294.08 | 0 | 8.28 |
| ENSMUSG00000058126 | Tpm3-rs7 | 0.20 | 44.44 | 0 | 7.66 |
| ENSMUSG00000063953 | Amd2 | 0 | 5.41 | 0 | 7.60 |
| ENSMUSG00000063696 | Gm8730 | 0.07 | 22.26 | 0 | 7.51 |
| ENSMUSG00000074516 | Gm10709 | 0.66 | 90.26 | 0 | 7.01 |
| ENSMUSG00000021908 | Gm6768 | 0.02 | 4.32 | 0 | 6.70 |
| ENSMUSG00000096403 | Gm9825 | 0.57 | 55.52 | 0 | 6.59 |
| ENSMUSG00000001348 | Acp5 | 4.33 | 0.15 | 0 | -4.70 |
| ENSMUSG00000073489 | Ifi204 | 0.97 | 0 | 2.63E-08 | -5.02 |
| ENSMUSG00000029307 | Dmp1 | 1.54 | 0.039 | 1.11E-16 | -5.09 |
| ENSMUSG00000039462 | Col10a1 | 0.75 | 0 | 2.10E-09 | -5.16 |
| ENSMUSG00000029306 | Ibsp | 5.75 | 0.13 | 0 | -5.38 |
| ENSMUSG00000050578 | Mmp13 | 3.90 | 0.04 | 0 | -6.16 |
| Gene ID | Gene Name | WT Lung | Dip2a<sup>-/-</sup> Lung | FDR | log2FC |
|  |  | FKPM | FKPM |  |  |
| ENSMUSG00000096403 | Gm9825 | 0.005 | 80.34 | 0 | 10.62 |
| ENSMUSG00000095427 | Rps2-ps6 | 0.17 | 30.4 | 0 | 7.12 |
| ENSMUSG00000074516 | Gm10709 | 0.25 | 34.49 | 0 | 6.65 |
| ENSMUSG00000074768 | Bhmt | 2.9 | 242.2 | 0 | 6.53 |
| ENSMUSG00000063696 | Gm8730 | 1.7 | 148.4 | 0 | 6.41 |
| ENSMUSG00000047676 | Rpsa-ps10 | 2.63 | 144.52 | 0 | 5.83 |
| ENSMUSG00000032315 | Cyp1a1 | 0.28 | 13.25 | 0 | 5.55 |
| ENSMUSG00000045027 | Prss22 | 0.55 | 23.05 | 0 | 5.32 |
| ENSMUSG00000078956 | Gm14221 | 5.32 | 1.12 | 1.01E-14 | 5.2 |
| ENSMUSG00000026073 | Il1r2 | 14.53 | 0.64 | 0 | -4.27 |
| ENSMUSG00000028341 | Nr4a3 | 24.47 | 0.99 | 0 | -4.48 |
| ENSMUSG00000023031 | Cela1 | 10.47 | 0.23 | 7.77E-16 | -4.68 |
| ENSMUSG00000047428 | Dlk2 | 3.84 | 0.086 | 1.01E-12 | -4.76 |
| ENSMUSG00000049796 | Crh | 6.74 | 0.13 | 1.11E-16 | -4.79 |
| ENSMUSG00000027313 | Chac1 | 10.42 | 0.2 | 0 | -5.23 |
| ENSMUSG00000020591 | Ntsr2 | 10.57 | 0.08 | 0 | -5.56 |

**Fig 2.**
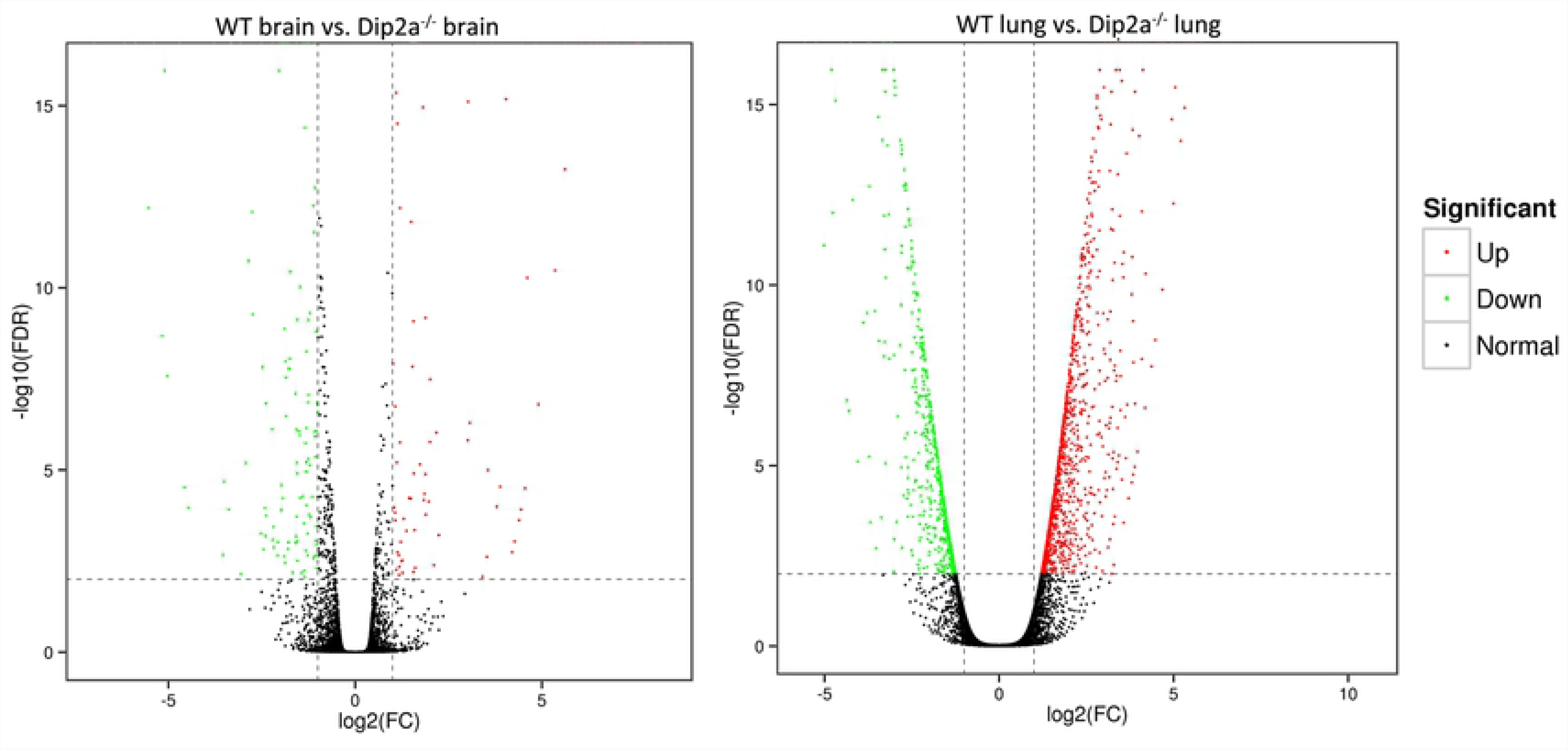
Differentially expressed genes volcano map. (a) WT brain vs. Dip2a^-/-^ brain. (b) WT lung vs. Dip2a^-/-^ lung. The red and green dots in the figure represent up-regulated and down-regulated differentially expressed genes respectively.

### GO enrichment analysis and COG classification of Dip2A-regulated DEGs

For gene ontology (GO) analysis, 185 DEGs from brain and 1709 DEGs from lung were classified into three GO categories and 51 terms (Fig 3). In biological process category, most of the DEGs in brain and lung were assigned to ‘cellular process’, ‘single-organism process’ and ‘metabolic process’. In molecular function category, most DEGs were annotated under ‘binding’, ‘catalytic activity’ and ‘signal transducer activity’. Within cellular component, ‘cell’, ‘cell part’ and ‘organelle’ was annotated with most DEGs. To further clarify the biological process, DEGs from both groups were enriched in 84 terms and the 10 most significant terms from each groups are summarized in figure 3 (c, d). In lung, the most significant biological terms include ‘regulation of transcription, DNA-templated’ and ‘positive-negative regulation of Transcription from RNA polymerase II promoter’ and ‘apoptotic process’. In brain, the most significant terms were ‘multicellular organism development’, ‘positive-negative regulation of Transcription from RNA polymerase II promoter’ and ‘cell differentiation’. In addition, 34 DEGs from lung and 12 DEGs from brain were annotated under GO term ‘in utero embryonic development’ (Fig 4). These DEGs are important in progression of embryo in uterus over time.

**Fig 3.**
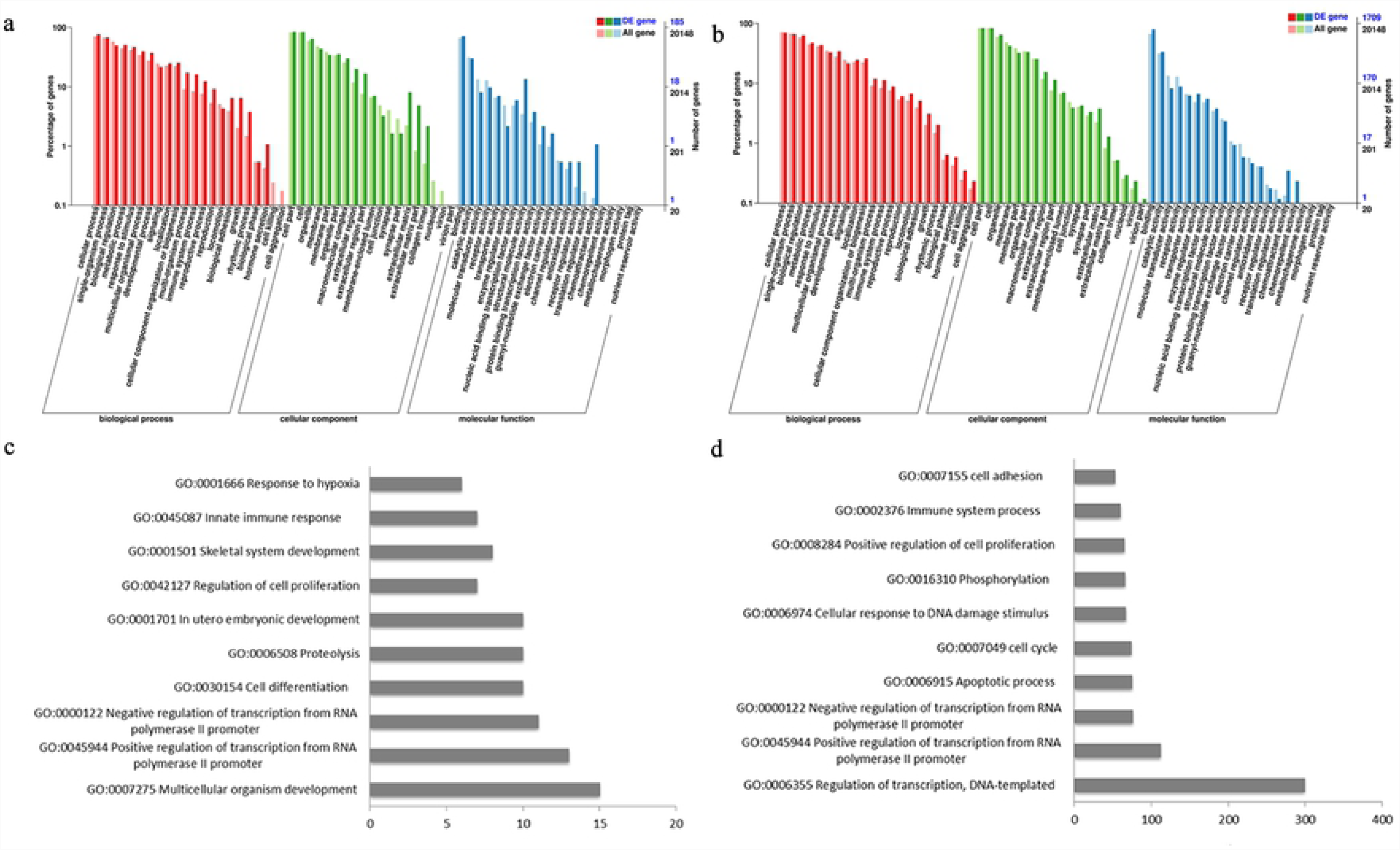
Gene Ontology (GO) classification of DEGs from WT brain vs. Dip2a^-/-^ brain (a, c) and WT lung vs. Dip2a^-/-^ lung (b, d). Histogram of GO annotation was generated by KOBAS (kobas.cbi.pku.edu.cn). The abscissa indicates GO classification, the Y-axis on the left indicates the percentage of genes, and the Y-axis on right indicates the number of genes. One unigenes could be assigned with more than one GO term. Most significant enriched biological terms in brain (c) and lung (d).

**Fig 4.**
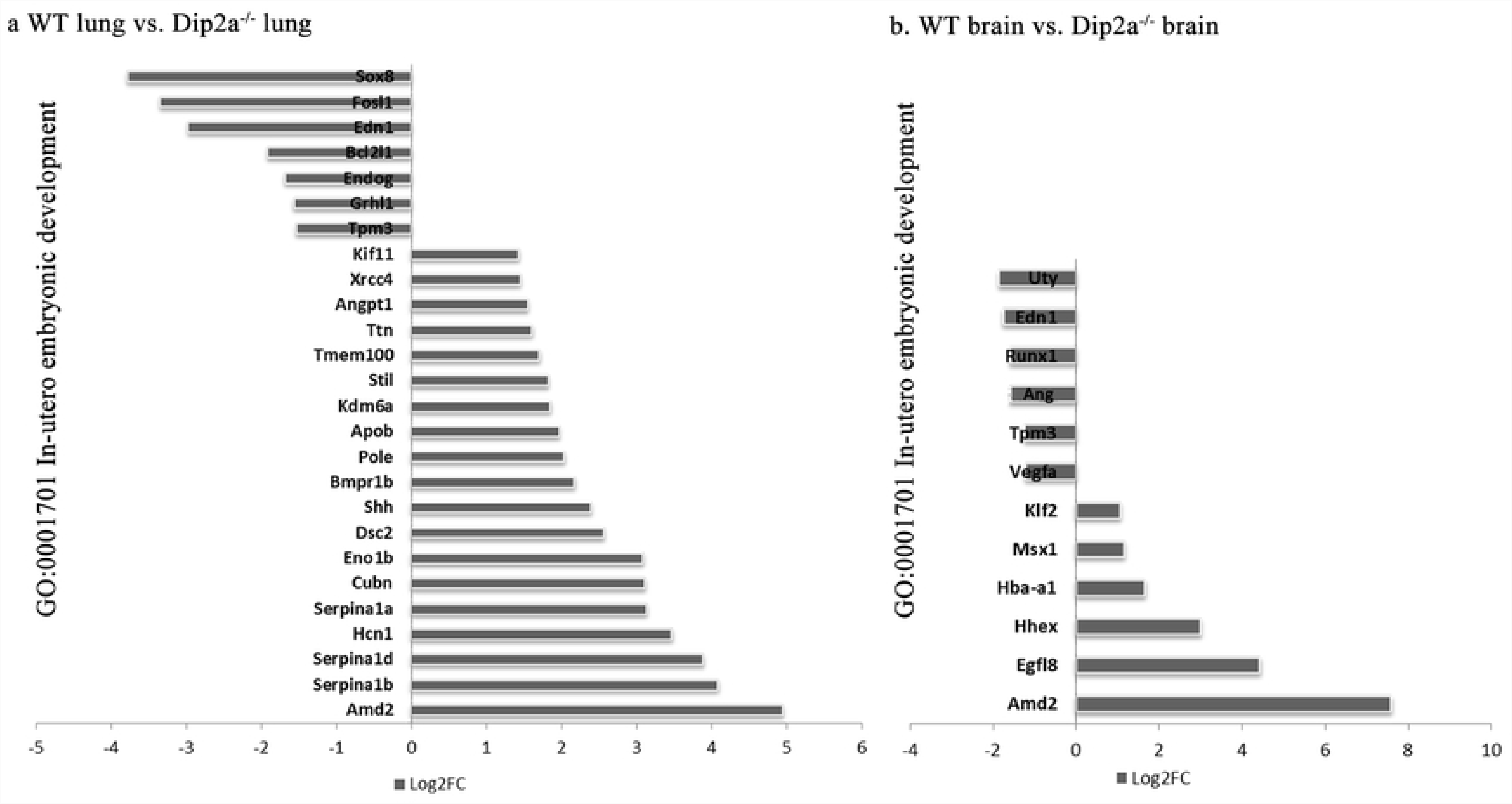
List of DEGs annotated to GO term ‘In-utero embryonic development’.

To further clarify the molecular function of Dip2A, total 54 and 677 DEGs from brain and lung were assigned to COG classification and divided into 26 specific categories (Fig 5). In both groups, the top hits include ‘Replication, Recombination and repair (7.25% & 10.86%)’, ‘Signal transduction and mechanism (5.8% and 8.69%)’, ‘Translation, Ribosomal structure and Biogenesis (2.61% &13.04%)’ and ‘Transcription (7.6% & 8.7%)’.

**Fig 5.**
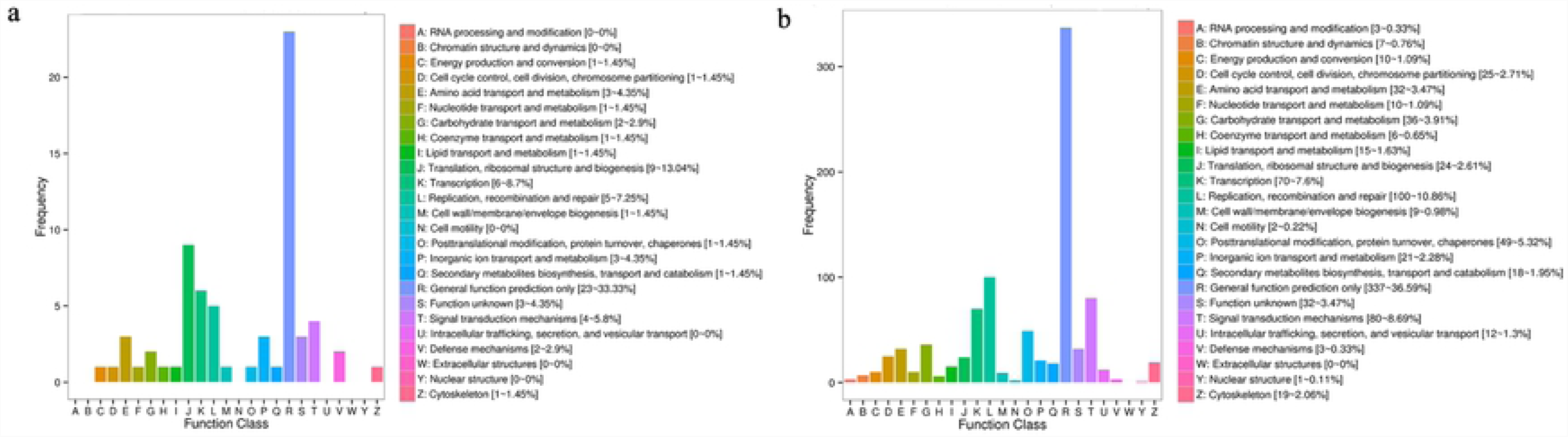
COG classification of differential expression genes (a) WT brain vs. Dip2a^-/-^ brain and (b) WT lung vs. Dip2a^-/-^ lung. The X-axix indicates content of each category of COG and the Y-axis indicates number of genes annotated in each category.

### KEGG pathway annotation of brain and lung DEGs

In the process of pathways annotation for Dip2A regulated DEGs, 70 DEGs from brain and 625 DEGs from lung were annotated to 112 and 264 pathways respectively in KEGG pathway database (S2 Fig). In order to analyze whether DEGs are over-presented on a pathway, the pathway enrichment analysis was performed (Fig 6). The top 5 enriched pathways in brain with the least significant Q value<0.05 and enrichment factor greater than 2 were ‘ko04610 Complement and coagulation cascades’, ‘ko05150 *Staphylococcus aureus* infection’, ‘ko01230 Biosynthesis of amino acids’, ‘ko04066 HIF-1 signaling pathway’ and ‘ko04151 PI3K-Akt signaling pathway’, whereas in lung, the most enriched pathways with the least Q value<1 and enrichment factor> 2 are ‘ko05216 Thyroid cancer’, ‘ko00740 Riboflavin metabolism’, ‘ECM-receptor interaction’, ‘ko05202 Transcriptional misregulation in cancer’ and ‘ko05200 Pathways in cancer’.

**Fig 6.**
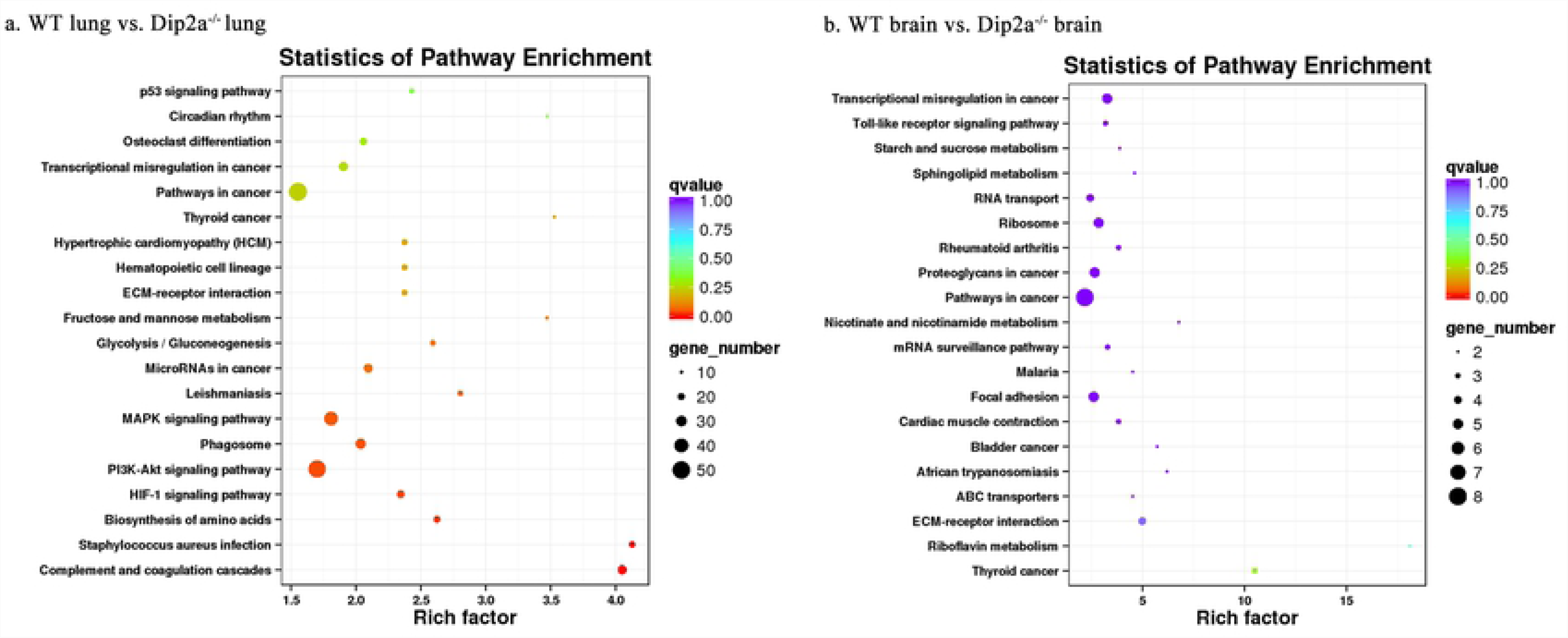
KEGG pathway enrichment scatter plot of DEGs. (a) DEGs of WT lung vs. Dip2a^-/-^ lung and (b) WT brain vs. Dip2a^-/-^ brain. Each circle in the figure represents a KEGG pathway. The Y-axis represents name of the pathway and the X-axis indicates Enrichment Factor, indicating the proportion of the annotated genes to all genes in the pathway.

### Transcription factor annotation of DEGs

Zinc finger domain containing transcription factor are the most abundant proteins whose function are extraordinarily diverse and include epithelium development, neo-cortex development, transcription activation, regulation of apoptosis, protein folding and assembly [18-19]. DIP2A is thought to be a transcription factor due to its zinc finger motif [2]. To extend these findings, 14 DEGs (9 up & 5 down) from brain and 203 DEGs (163 up & 40 down) from lung were annotated with transcription factor (animal TFDB) database. In both group, the most of up-regulated genes belongs to Zinc finger Cys_2_His_2_-like class group (ZF-C2H2) [124 & 2], Homeobox (5 & 2), High-mobility group (HMG) [4 & 1], Zinc finger and BTB domain-containing protein (ZBTB) [4 & 1], whereas the most of down-regulated genes accounts to transcription factor basic leucine zipper domain (TF-bZIP) [8 & 1], Thyroid hormone receptor [2&1] and Interferon regulatory factor (IRF) [2,1]. Based upon these evidences, our study strongly suggests that Dip2a protein regulate expression of Zinc Finger domain containing proteins during lung and brain development. Transcription factor with the highest fold change (FC>6) from each group is listed in Table 3.

**Table 3.** List of highly differentially expressed Transcription factors (FC>6, FDR<0.001) in WT lung vs. Dip2a^-/-^ lung and WT brain vs. Dip2a^-/-^ brain respectively.

| Gene ID | Gene Symbol | log2FC | TF Family |
| --- | --- | --- | --- |
| ENSMUSG00000090093 | Gm14399 | 4.118479 | zf-C2H2 |
| ENSMUSG00000056824 | Zfp663 | 3.892387 | zf-C2H2 |
| ENSMUSG00000074867 | Zfp808 | 3.841335 | zf-C2H2 |
| ENSMUSG00000078902 | Gm14443 | 3.669152 | zf-C2H2 |
| ENSMUSG00000078864 | Gm14322 | 3.530807 | zf-C2H2 |
| ENSMUSG00000031079 | Zfp300 | 3.363612 | zf-C2H2 |
| ENSMUSG00000074865 | Zfp934 | 3.350508 | zf-C2H2 |
| ENSMUSG00000078502 | Gm13212 | 3.262812 | zf-C2H2 |
| ENSMUSG00000061371 | Zfp873 | 3.252192 | zf-C2H2 |
| ENSMUSG00000090015 | Gm15446 | 3.222472 | zf-C2H2 |
| ENSMUSG00000046351 | Zfp322a | 3.212515 | zf-C2H2 |
| ENSMUSG00000055240 | Zfp101 | 3.180478 | zf-C2H2 |
| ENSMUSG00000030393 | Zik1 | 3.146957 | zf-C2H2 |
| ENSMUSG00000069184 | Zfp72 | 3.037894 | zf-C2H2 |
| ENSMUSG00000078899 | Gm4631 | 3.009177 | zf-C2H2 |

| ENSMUSG00000028341 | Nr4a3 | -4.48349 | NOR |
| --- | --- | --- | --- |
| ENSMUSG00000024176 | Sox8 | -3.77856 | HMG |
| ENSMUSG00000024912 | Fosl1 | -3.34232 | TF_bZIP |
| ENSMUSG00000003545 | Fosb | -3.01894 | TF_bZIP |
| Gene ID | Gene Symbol | log2FC | TF Family |
| ENSMUSG00000001444 | Tbx21 | 4.909944 | T-box |
| ENSMUSG000000067261 | Foxd3 | 4.204621 | Fork head |
| ENSMUSG000000055102 | Zfp819 | 3.790954 | zf-C2H2 |
| ENSMUSG000000022479 | Vdr | -3.50616 | Thyroid receptor<br>hormone |
| ENSMUSG000000070031 | Sp140 | -3.024071 | SAND |
| ENSMUSG000000024986 | Hhex | -3.013431 | Homeobox |

### DEGs validation by quantitative real-time PCR

To evaluate validity of RNA-Seq data, five up-regulated DEGs and five-down regulated DEGs from each group were selected for quantitative real-time RT-PCR (qPCR) (Fig 7). The RNA-Seq results of these genes were similar to those obtained by qPCR. These results confirmed the good quality of RNA-Seq results.

**Fig 7.**
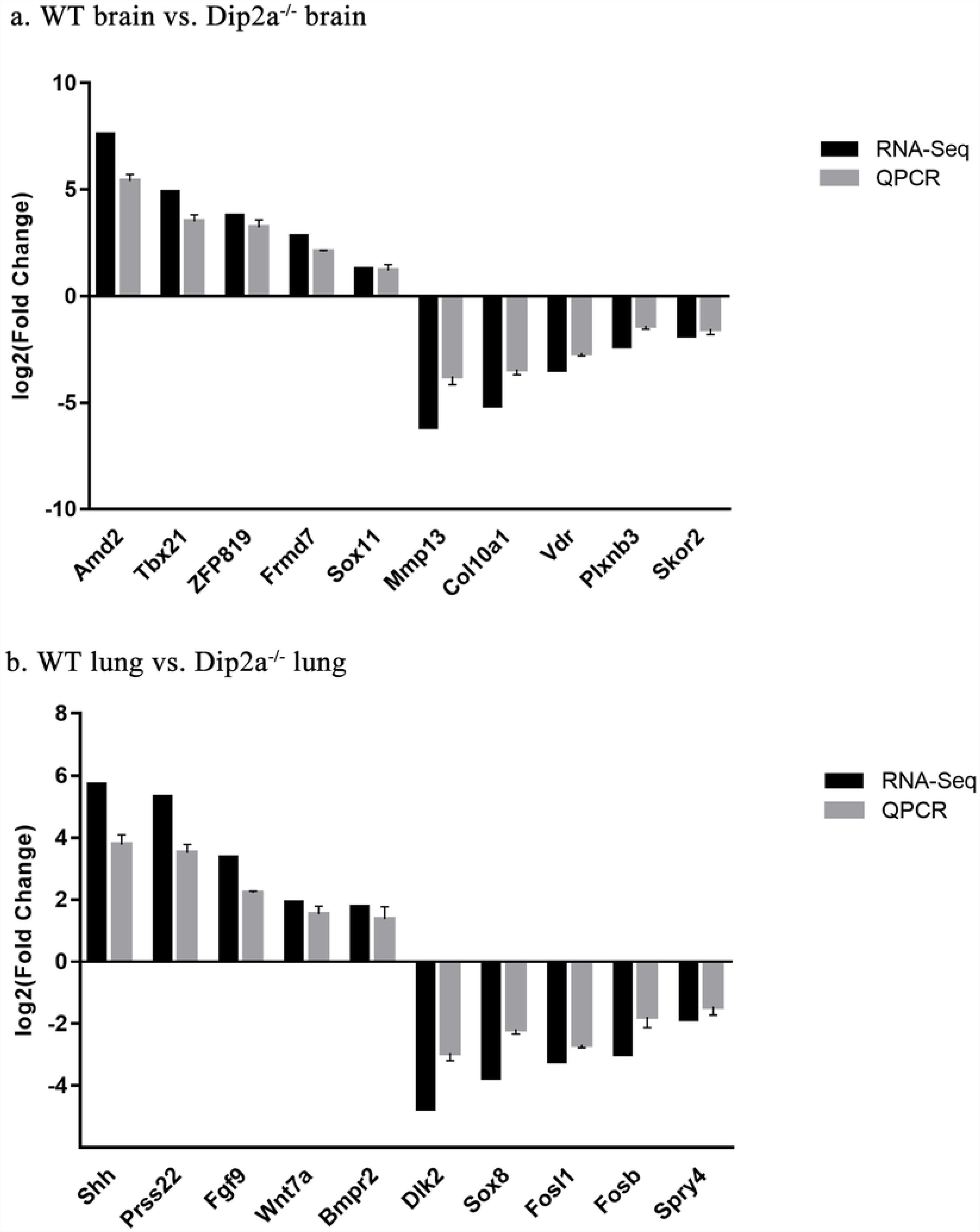
Validation of RNA-Seq results by real-time quantitative PCR (QPCR).

### Roles of Dip2A in neuronal cell maturation, differentiation and survival

Previous studies have suggested that Dip2a is highly expressed in neuronal cells of developing central nervous system such as retinal ganglion cells, Purkinje cell layer and granular cell, and may play important roles in synapse formation and axon guidance [1-4]. In this study, we found 10 genes that are important in neuronal cell maturation and in brain development were differentially expressed in Dip2a^-/-^ brain. Skor2 and Gpr37l1 genes important in Purkinje cell maturation, differentiation and layer formation were down-regulated [20-21]. Runx1 gene is an important in cell fate specification and axonal projections of dorsal root ganglion neurons and Erbb3 gene is required in the control of growth and development of Schwann cell. These genes were down-regulated [22-23]. Similarly, Frmd7 gene which promotes neuronal outgrowth and migration of neural precursor cell was up-regulated [24]. Fut10 is important in maintenance and differentiation of neuron stem cell and was up-regulated [25]. Extracellular matrix component Hapln1 gene that plays an important in neo-cortex development and expansion was found over expressed [26]. Transcription factor SRY-box (Sox) family gene Sox11 is expressed abundantly in all type of embryonic sensory neurons including sensory ganglion and trigeminal ganglion and promotes neuronal maturation was found up-regulated [27]. In addition, transcription factor AP-2 family gene Tfap2c important in neural crest induction was under expressed [28]. We also found Plxnb3 gene was down-regulated. Increasing evidence suggests that Plexin-B3 is axon guidance molecule and promotes synapse formation in rat hippocampal neurons [29]. Hence, these finding strongly supports the role of Dip2a in all type of neuronal cell maturation, differentiation and survival.

### Roles of Dip2A in lung morphogenesis and development

Lung development has been divided into four stages in both human and mice. From E17.5 to postnatal day 5 (P5) in mouse is a terminal saccular stage where substantial thinning of interstitium occurs as a result of apoptosis as well as differentiation of mesenchyme cells [30]. In this study, 132 DEGs from lung were annotated under biological term apoptosis process (Fig 3d). Up-regulated genes are Ddias, Terf1, Tnfsf10, Stk26 and Thbs2 (Log2FC> 3) and down-regulated genes include Gzmb, Bcl2l1, Thbs1 and Gzma (Log2FC> −3). In addition, we also found Hoxa5, Eya1, Ctsh, Erff1, Lama1, Lama2, Rspo2, Sox11, Spry4, Shh, Igf1 and Wnt7a, important in lung epithelial, mesenchyme, vascular development and branching morphogenesis, were over expressed [31-40]. Taking together, this study supports the roles of Dip2A in apoptosis and organogenesis during embryonic lung development.

## Conclusion

In this report, four Transcriptome, including WT brain and lung, Dip2a^-/-^ brain and lung at embryonic E19.5 were analyzed. On an average 6000 unigenes in each sample were generated with the Illumina HiSeq 2500 platform. In WT brain vs. Dip2a^-/-^ brain comparison, a total of 214 DEGs were detected, including 82 up- and 132 down-regulated genes. These DEGs included genes involved in neuronal cell maturation, differentiation and survival. In WT lung vs. Dip2a^-/-^ lung comparison, a total of 1900 DEGs were detected, including 1259 up- and 641 down-regulated genes. These DEGs are important in apoptosis process, lung epithelial development and in morphogenesis. To conclude, we have identified several candidate genes that are regulated by Dip2A at E19.5 brain and lung. It would be interesting to further study the biological functions of these genes in brain and lung development.

## Acknowledgments

We are very thankful to Huiyan Wu and Xiu lu for microinjection and mouse colony management.

## Supporting information

**S1 Table. BLAST analysis of the non-redundant DEGs against six public databases.**

**S2 Fig. Annotated diagram of the KEGG pathway of differentially expressed genes; (a) WT lung vs. Dip2a**^**-/-**^ **lung (b) WT brain vs. Dip2a**^**-/-**^ **brain.**

## References

1. Steller H, Fischbach KF & Rubin GM. Disconnected: a locus required for neuronal pathway formation in the visual system of Drosophila. Cell. 1987; 50:1139–1153.

2. Heilig JS, Freeman M, Laverty T, Lee KJ, Campos AR, Rubin GM & Steller H. Isolation and characterization of the disconnected gene of Drosophila melanogaster. EMBO J. 1991; 10:809–815.

3. Campos AR, Lee KJ & Steller H. Establishment of neuronal connectivity during development of the Drosophila larval visual system. J Neurobiol. 1995; 28:313–329.

4. Mukhopadhyay M, Pelka P, DeSousa D, Kablar B, Schindler A, Rudnicki MA, et al. Cloning, genomic organization and expression pattern of a novel Drosophila gene, the disco-interacting protein 2 (dip2), and its murine homolog. Gene. 2002; 293(1–2):59–65. PMID: 12137943.

5. Tanaka M, Murakami K, Ozaki S, Imura Y, Tong XP, Watanabe T, et al. DIP2 disco-interacting protein 2 homolog A (Drosophila) is a candidate receptor for follistatin-related protein /follistatin-like1—analysis of their binding with TGF-beta super family proteins. FEBS J. 2010; 277(20):4278–89.doi:10.1111/j. 1742-4658. PMID:20860622.

6. Guan JS, Haggarty SJ, Giacometti E, Dannenberg JH, Joseph N, Gao J et al. HDAC2 negatively regulates memory formation and synaptic plasticity. Nature. 2009; 459(7243):55–60. doi:10.1038/nature07925.

7. Egger, G., Roetzer, K.M., Noor, A. et al. Identification of risk genes for autism spectrum disorder through copy number variation analysis in Austrian families. Neurogenetics. 2014; 15:117. doi:10.1007/s10048-014-0394-0.

8. De Jager, P., G. Srivastava, K. Lunnon, J. Burgess, L. Schalkwyk, L. Yu, M. Eaton, et al. Alzheimery’s disease pathology is associated with early alterations in brain DNA methylation at ANK1, BIN1, RHBDF2 and other loci. Nature neuroscience. 2014; 17 (9): 1156–1163. doi:10.1038/nn.3786.

9. Zhang L, Jia R, Palange NJ, Satheka AC, Togo J, An Y, et al. Large genomic fragment deletions and insertions in mouse using CRISPR/Cas9. PLoS One. 2015; 10(3):e0120396. doi:10.1371/journal.pone.0120396 PMID: 25803037.

10. Zhang L, Mabwi HA, Palange NJ, Jia R, Ma J, Bah FB, et al. Expression Patterns and Potential Biological Roles of Dip2a. PLoS One. 2015; 10 (11): e0143284. doi:10.1371 /journal pone. 0143284

11. Ozsolak F. & Milos, P. M. RNA sequencing: advances, challenges and opportunities. Nature Reviews Genetics. 2015; 12(2), 87–98. doi:10.1038/nrg2934.

12. Trapnell C, Roberts A, Goff L, Pertea G, Kim D, Kelley DR, et al. Differential gene and transcript expression analysis of RNA-seq experiments with TopHat and Cufflinks. Nat Protoc. 2012; 7:562–578. doi:10.1038/nprot.2012.016. PMID: 22383036

13. Altschul SF, Madden TL, Schaffer AA, et al. Gapped BLAST and PSI-BLAST: a new generation of protein database search programs. Nucleic Acids Res. 1997; 25:3389–3402.

14. Conesa A, Gotz S, García-Gómez JM, Terol J, Talón M, Robles M. Blast2GO: a universal tool for annotation, visualization and analysis in functional genomics research. Bioinformatics. 2005; 21(18):3674–3676 PMID:16081474.

15. Anders S, Huber W. Differential expression analysis for sequence count data. Genome Biology. 2010; doi:10.1186/gb-2010-11-10-r106.

16. Young MD, Wakefield MJ, Smyth GK, et al. Gene ontology analysis for RNA-seq: accounting for selection bias. Genome Biology. 2010; doi:10.1186/gb-2010-11-2-r14.

17. Hong Li, Meifang Lu, Liu X. Zinc-Finger Proteins in Brain Development and Mental Illness. J Transl Neurosci. 2018; 3:4. doi:10.21767/2573-5349.100017.

18. John H Laity, Brian M Lee, Peter E Wright, Zinc finger proteins: new insights into structural and functional diversity. Current Opinion in Structural Biology. 2001; 11(1). doi.:10.1016/S0959-440X (00)00167-6.

19. Baiping Wang, Wilbur R. Harrison, Hui Zheng. Transposon mutagenesis with coat color genotyping identifies an essential role for Skor2 in sonic hedgehog signaling and cerebellum development. Development. 2011; 138(20):4487–4497. doi:10.1242/dev.067264. PMID:21937600

20. Marazziti, Daniela et al. “Precocious cerebellum development and improved motor functions in mice lacking the astrocyte cilium-, patched 1-associated Gpr37l1 receptor.” Proceedings of the National Academy of Sciences. 2013; 110(41):16486–16491. doi:10.1073/pnas.1314819110.

21. Masaaki Yoshikawa, Kouji Senzaki, Tomomasa Yokomizo, Satoru Takahashi, Shigeru Ozaki, Takashi Shiga. Runx1 selectively regulates cell fate specification and axonal projections of dorsal root ganglion neurons. Developmental Biology. 2007; 303(2); 663–647. doi:10.1016/j.ydbio.2006.12.007.

22. Stefan Britsch, Li, Susanne Kirchhoff, Franz Theuring, et. al. The ErbB2 and ErbB3 receptors and their ligand, neuregulin-1, are essential for development of the sympathetic nervous?system. Genes & Dev. 1998. 12:1825–1836. doi:10.1101/gad.12.12.1825.

23. Betts-Henderson JL, Bartesaghi S, Crosier M, Lindsay S, et. al. The nystagmus-associated FRMD7 gene regulates neuronal outgrowth and development. Hum Mol Genet. 2010; 19(2):342–51. doi:10.1093/hmg/ddp500.

24. Akhilesh Kumar, Tomohiro Torii, Yugo Ishino, Daisuke Muraoka, et. al. The Lewis X-related α1,3-Fucosyltransferase, Fut10, Is Required for the Maintenance of Stem Cell Populations. J Biol Chem. 2013; 288(40):28859–28868. doi:10.1074/jbc.M113.469403.

25. Katherine R. Long, Ben New land, Marta Florio, Nereo Kalebic, et. al. Extracellular Matrix Components HAPLN1, Lumican, and Collagen I Cause Hyaluronic Acid-Dependent Folding of the Developing Human Neocortex. 2018; 99(4):702–719. doi.10.1016/j.neuron.2018.07.013.

26. Lin, L., Lee, V., Wang, Y., Lin, J., Sock, E., Wegner, M. and Lei, L. Sox11 regulates survival and axonal growth of embryonic sensory neurons. Dev. Dyn. 2011; 240:52–64. doi:10.1002/dvdy.22489.

27. D.J. Coelho, D.J. Sims, P.J. Ruegg, I. Minn, A.R. Muench, P.J. Mitchell, Cell type-specific and sexually dimorphic expression of transcription factor AP-2 in the adult mouse brain. Neuroscience. 2005; 134(3):907–919. doi:10.1016/j.neuroscience.2005.04.060.

28. Piret Laht, Epp Tammaru, Maarja Otsus, Johan Rohtla, Liivi Tiismus, Andres Veske. Plexin-B3 suppresses excitatory and promotes inhibitory synapse formation in rat hippocampal neurons. Experimental Cell Research. 2015; 335(2):269–278. doi:10.1016/j.yexcr.2015.05.007.

29. David Warburton, Ahmed El-Hashash, Gianni Carraro, Caterina Tiozzo, et. al. Curr Top Dev Biol. 2010; 90:73–158. doi:10.1016/S0070-2153(10)90003-3.

30. Claudia Kappen. Hox Genes in the Lung. Am J Respir Cell Mol Biol. 1996; 15(2):156–162. doi:10.1165/ajrcmb.15.2.8703471.

31. El-Hashash AH1, Al Alam D, Turcatel G, Bellusci S, Warburton D. Eyes absent 1 (Eya1) is a critical coordinator of epithelial, mesenchymal and vascular morphogenesis in the mammalian lung. Dev Biol. 2011; 350(1):112–26. doi:10.1016/j.ydbio.2010.11.022.

32. Jining Lü, Jun Qian, Daniel Keppler, Wellington V. Cardoso. Cathespin H Is an Fgf10 Target Involved in Bmp4 Degradation during Lung Branching Morphogenesis. 2007; 282:22176–22184. doi:10.1074/jbc.M700063200.

33. Jin N, Cho SN, Raso MG, Wistuba II, Smith Y, Yang Y et al. Mig-6 is required for appropriate lung development and to ensure normal adult lung homeostasis. Development. 2009; 136(19):3347–3356. doi:10.1242/dev.03297.

34. Schuger L, Skubitz AP, Zhang J, Sorokin L, He L. Laminin alpha1 chain synthesis in the mouse developing lung: requirement for epithelial-mesenchymal contact and possible role in bronchial smooth muscle development. J Cell Biol. 1997; 139(2):553–62.

35. Bell, Sheila M. et al. R-spondin 2 is required for normal laryngeal-tracheal, lung and limb morphogenesis.” Development. 2008; 135(6):1049–1058. doi:10.1242/dev.013359.

36. Zhu Y, Li Y, Jun Wei JW, Liu X. The role of Sox genes in lung morphogenesis and cancer. Int J Mol Sci. 2012; 13(12):15767–83. doi:10.3390/ijms131215767.

37. Taniguchi K, Ayada T, Ichiyama K, Kohno R, Yonemitsu Y, Minami Y, Kikuchi A, Maehara Y, Yoshimura A. Sprouty2 and Sprouty4 are essential for embryonic morphogenesis and regulation of FGF signaling. Biochem Biophys Res Commun. 2007; 352(4):896–902. doi:10.1016/j.bbrc.2006.11.107.

38. Van Tuyl M, Post M. From fruitflies to mammals: mechanisms of signalling via the sonic hedgehog pathway in lung development. Respir Res. 2000;1(1):30–5. doi:10.1186/rr9.

39. Zheng Wang, Wenting Li, Qiongya Guo, Yuming Wang, Lijun Ma, Xiaoju Zhang. Insulin-Like Growth Factor-1 Signaling in Lung Development and Inflammatory Lung Diseases. 2018; ID 6057589. https://doi.org/10.1155/2018/6057589.

40. Donna R. Newman 1, Huiying Zhang, Katherine Bortoff, James C. Bonner, and Philip L. Sannes. Alveolar Epithelial Differentiation during Repair Involves FoxA1, Wnt7A, and TGF-b. 2010; 7(2).

